# A mean-field to capture asynchronous irregular dynamics of conductance-based networks of adaptive quadratic integrate-and-fire neuron models

**DOI:** 10.1101/2023.06.22.546071

**Authors:** Christoffer G. Alexandersen, Chloé Duprat, Aitakin Ezzati, Pierre Houzelstein, Ambre Ledoux, Yuhong Liu, Sandra Saghir, Alain Destexhe, Federico Tesler, Damien Depannemaecker

**Affiliations:** Mathematical Institute, University of Oxford, Oxford, UK; Paris Saclay University, Institute of Neuroscience, CNRS, Saclay, France; Institut de Neurosciences des Systèmes, Aix-Marseille University, INSERM, Marseille, France; Group for Neural Theory, LNC2, INSERM U960, DEC, École Normale Supérieure - PSL University, Paris, France; Institute of Physiological Chemistry, Johannes Gutenberg University of Mainz, Mainz, Germany; Institute of Experimental Epileptology and Cognition Research, University of Bonn Medical Center, Bonn, Germany; Department of Software Engineering and Theoretical Computer Science, Technische Universität Berlin, Berlin, Germany

## Abstract

Mean-field models are a class of models used in computational neuroscience to study the behaviour of large populations of neurons. These models are based on the idea of representing the activity of a large number of neurons as the average behaviour of “mean field” variables. This abstraction allows the study of large-scale neural dynamics in a computationally efficient and mathematically tractable manner. One of these methods, based on a semi-analytical approach, has previously been applied to different types of single-neuron models, but never to models based on a quadratic form. In this work, we adapted this method to quadratic integrate-and-fire neuron models with adaptation and conductance-based synaptic interactions. We validated the mean-field model by comparing it to the spiking network model. This mean-field model should be useful to model large-scale activity based on quadratic neurons interacting with conductance-based synapses.

## 1 Introduction

Modelling brain activities over different scales is currently is a very relevant challenge. Many models were built over the years, going from sub-cellular to whole-brain scales, to serve various purposes and applications. In order to model the mesoscopic scale in particular, one option is to build phenomenological neural-mass models describing observations made at this scale. Another alternative consists in a bottom-up approach where the dynamics of the mesoscopic scale are derived by developing a mean-field model of the microscopic scale, i.e. of spiking neural network models. The mean-field approximation is a powerful tool for modelling the behaviour of large populations of neurons, enabling multiple applications [1]. In the last decade many mean-field approaches based on different spiking models have been proposed [2–9]. One of these methods, based on a semi-analytical approach [2, 4], was successfully applied to many single-neuron models [3, 5], but never to a quadratic neuron model. In this work, we aimed to adapt and apply this method to a quadratic neuron model proposed by Izhikevich [10].

Mean-field or neural-mass models are appropriate to model clinically recorded signals such as fMRI, EEG, or MEG [11, 12] because they provide a simplified representation of the complex electrical and synaptic activity of large populations of neurons. These models are based on the idea that the activity of a large group of neurons can be described by the average electrical activity of the group, without having to consider the individual activity of each neuron.

One of the most widely used models in computational neuroscience is the quadratic integrate-and-fire neuron model (QIF). In this model, the membrane potential of a single neuron is described by a quadratic differential equation, and spikes are generated when the membrane potential reaches a given threshold. In an extension of this model, called the adaptive-quadratic-integrate-and-fire neuron model (aQIF), a second slower variable describes the adaptive behaviour and enables the system to capture a large repertoire of electrophysiological patterns [10].

In this work, we build a model of cortical columns based on a balanced spiking network of aQIF neurons, composed of excitatory regular-spiking and inhibitory fast-spiking populations interacting through conductance-based synapses. This sparse network exhibits asynchronous irregular dynamics [13] as observed in awake brain states. We build the corresponding mean-field based on a previously developed master-equation formalism [2, 3, 14] and we compare its dynamics to the spiking neural network. We show that the mean-field can correctly capture the dynamics of the network for both constant and time-varying inputs. In addition, we show that our semi-analytical model remains valid for a wide range of cell parameters which guarantees the robustness and generality of this formalism.

## 2 Methods

### 2.1 Spiking network model

To build consider the spiking network model we consider the point neuron model proposed by Izhikevich [10] (Equations (1) & (2)).

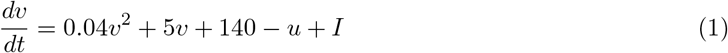

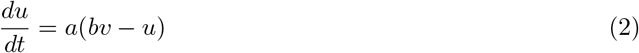

These equations can be rewritten in the following form.

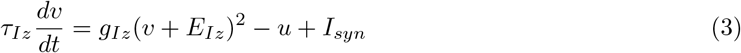

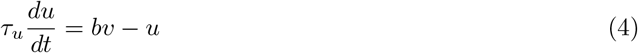

When an action potential is emitted (i.e. the membrane potential crosses a threshold), the system is reset as in the Equation (5):

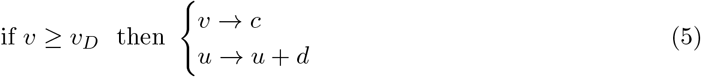

With *c* = *−*60, *d* = 0,*g*_*Iz*_ = 0.04 and *E*_*Iz*_ = *−*60 for FS cells, and, *c* = *−*65, *d* = 15, *g*_*Iz*_ = 0.01 and *E*_*Iz*_ = − 65 for RS cell, with *V*_*D*_ = − 30, and *τ*_*u*_ = 1 for both populations. Then, neuronal interactions are mediated through synaptic currents (Equation (6)).

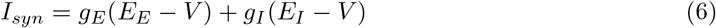

where *g*_*E,I*_ is the conductance of the excitatory and inhibitory synapses respectively, and *E*_*E,I*_ is the corresponding reversal potential. We model the conductances *g*_*E,I*_ as a decaying exponential function that takes kicks of amount *Q*_*E,I*_ at each presynaptic spike:

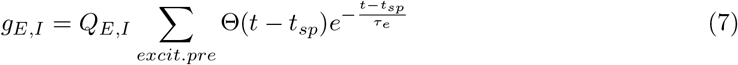

where the sum goes through all presynaptic excitatory spikes, Θ is the heaviside function, *τ*_*e*_ = *τ*_*i*_ = 5 ms is the decay timescale of excitatory and inhibitory synapses, and *Q*_*E*_ = 1.5 nS (*Q*_*I*_ = 5 nS) the excitatory (inhibitory) quantal conductance (i.e. the change in conductance generated by a single spike).

### 2.2 Mean-field model

To build the mean-field model of our system, we follow the formalism proposed by El Boustani and Destexhe [2, 3]. This formalism provides a second-order mean-field allowing us to derive a set of differential equations that describe the evolution of the mean firing rate *v*_*µ*_ of each population (Equation (8)), the covariance 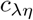 between populations *λ/η* (Equation (9)), and the average adaptation for the excitatory population *u* (Equation (10)), with *µ, λ, η* = *e, i*.

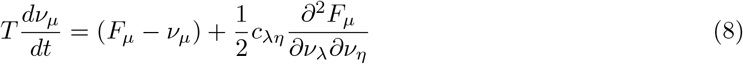

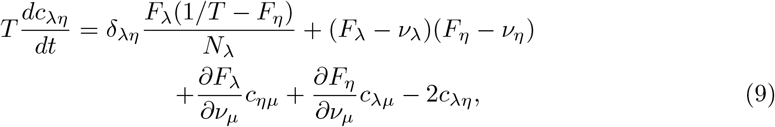

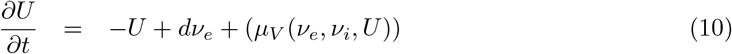

The parameter *T*, the time constant of the firing rate equations and covariance equations, is related with a main assumption used in this mean-field derivation: the network dynamics is considered to be markovian within a time resolution T [2] [2]. It is thus assumed that each neuron emits a maximum of one single spike in a time window of length T. As such, the range of firings rates that are valid under the assumptions of the mean-field are constrained by the value of *T* .

The functions *F*_*e*_ and *F*_*i*_ correspond respectively to the transfer function of the excitatory and inhibitory neurons (i.e. each neural subtype’s output firing rate when receiving excitatory and inhibitory inputs with rates *v*_*e*_ and *v*_*i*_). They are a function of the firing rates and of the adaptation: *F*_*e,i*_(*v*_*e*_ + *v*_*ext*_, *v*_*i*_, *U*), where *v*_*ext*_ is the firing rate of an external drive, corresponding to the Poissonian external input in the spiking network model. To build this transfer function, we first need to numerically calculate the output firing rate of the single neuron model for varying excitatory and inhibitory inputs. To account for the nonlinearity of the dynamics of the single neuron model (spike generation and reset), we consider an effective or phenomenological threshold, 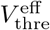, a function of the statistics of the subthreshold membrane voltage dynamics. This statistic is assumed to be normally distributed, with the average membrane voltage *µ*_*V*_, its standard deviation σ_*V*_ and auto-correlation time *τ*_*V*_, their calculation are described below.

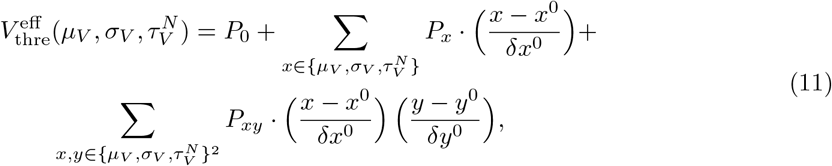

The next step within our semi-analytic approach is to fit the output firing rate of the single neuron with the template transfer function given in Equation (12), where *erfc* is the Gauss error function.

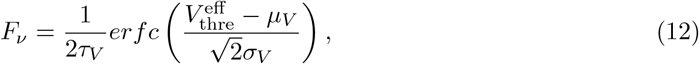

Considering asynchronous irregular regimes [13], we make the assumption that the input spike trains follow a Poissonian statistics [2–4] to calculate the averages (*µ*_*Ge,Gi*_) and standard deviations (*σ*_*Ge,Gi*_) of the conductances, described in Equations 13. In these equations, *K*_*e*_ and *K*_*i*_ are the average input connectivity received from the excitatory or inhibitory population respectively. Like in the spiking network, *τ*_*e*_ = *τ*_*i*_ = *τ*_*syn*_ are synaptic time constants and *Q*_*e*_ and *Q*_*i*_ are the quantal increments of the conductances respectively for the excitatory or inhibitory population.

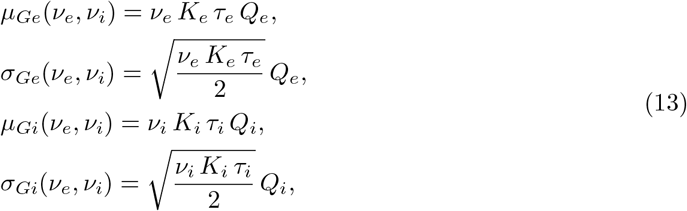

Then we can calculate the mean subthreshold voltage by taking the stationary solution of Equation 3. The quadratic form of Equation 3 in *v* gives rise to two solutions, of which one is identified as diverging and, as such, discounted. We thus obtain Equation 14, which differs from the form obtained for other point neuron models [3–5]. Then, applying the approach described in previous work [4], we determine *σ*_*v*_ and *τ*_*v*_ as follows:

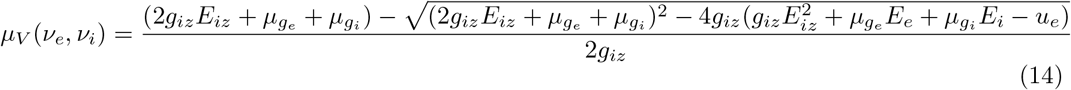

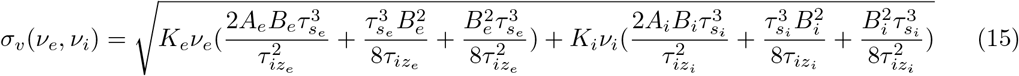

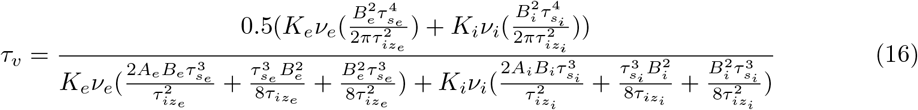

Where

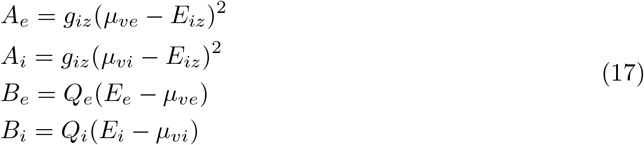

## 3 Results

In this section we will implement and validate the mean-field model for the quadratic Izhikevich model [10] described in the previous sections. The first step to implement the mean-field formalism consists in the estimation of the semi-analytical transfer function (TF) described in Eqs. 11 and 12. The parameters of the TF are estimated by fitting the template transfer function with the output firing rate obtained numerically from a single Izhikevich neuron for varying inputs rate *v*_*e*_ and *v*_*i*_. The output of the single neuron is shown in Fig. 1.a. In Fig. 1.d, we show the comparison of the semi-analytical TF obtained from the fit with the corresponding output rate of the single neuron. We see that the TF accurately captures the output rate obtained from the numerical simulations.

**Figure 1.**
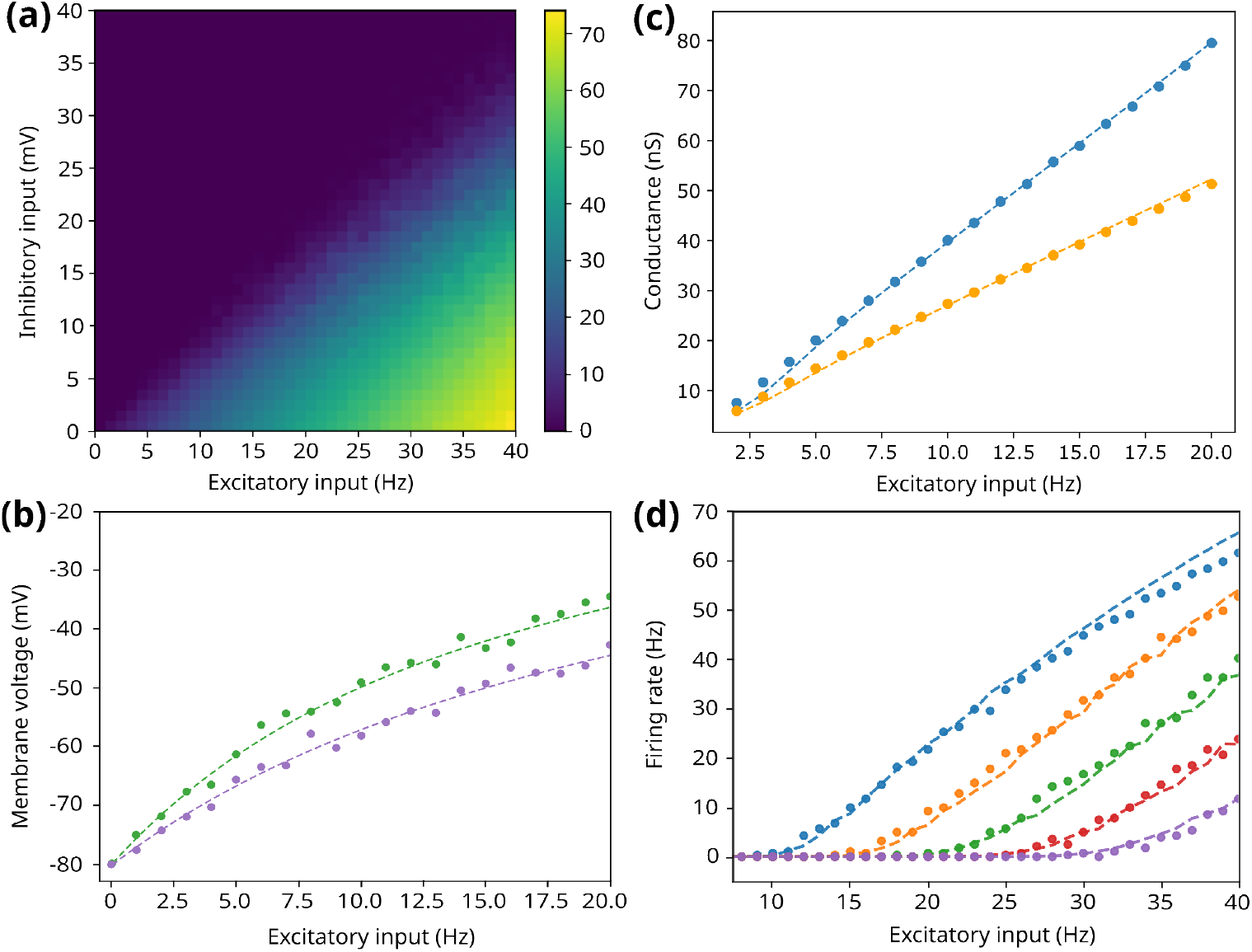
: Validation of the transfer function used for the mean-field formalism. a) Numerical transfer function obtained from a single aQIF cell. b-c) Mean excitatory (b, blue) and inhibitory (b, yellow) conductances and mean membrane potential (c, different colors for different inhibitory inputs) obtained from numerical simulations together with the corresponding prediction from the mean-field. d) Firing rates obtained from single cell simulations together with the prediction from the semi-analytical transfer function. Each curve corresponds to different inhibitory inputs.

Two other relevant quantities predicted from the mean-field formalism are the membrane potential and the synaptic conductances. We display in Fig.1.b and c the values obtained numerically for these quantities together with the mean-field prediction as a function of the excitatory input rate, showing a good match between the numerical value and the one expected from the mean-field.

### 3.1 Spontaneous activity and second order MF evaluation

Once the transfer function has been obtained and validated, we continue with the analysis of the mean-field response and its comparison with the corresponding network simulations. We begin with analyzing the response of the mean-field to constant external excitatory drives. We show in Fig. 2.a the results of the firing rates of the mean-field together with the results obtained from the network as a function of the external excitatory drive. In addition, we show in Figs. 2.b and c the distribution of firing rates obtained from the network together with the distribution predicted by the second order mean-field (Eqs.8 and 9).

**Figure 2.**
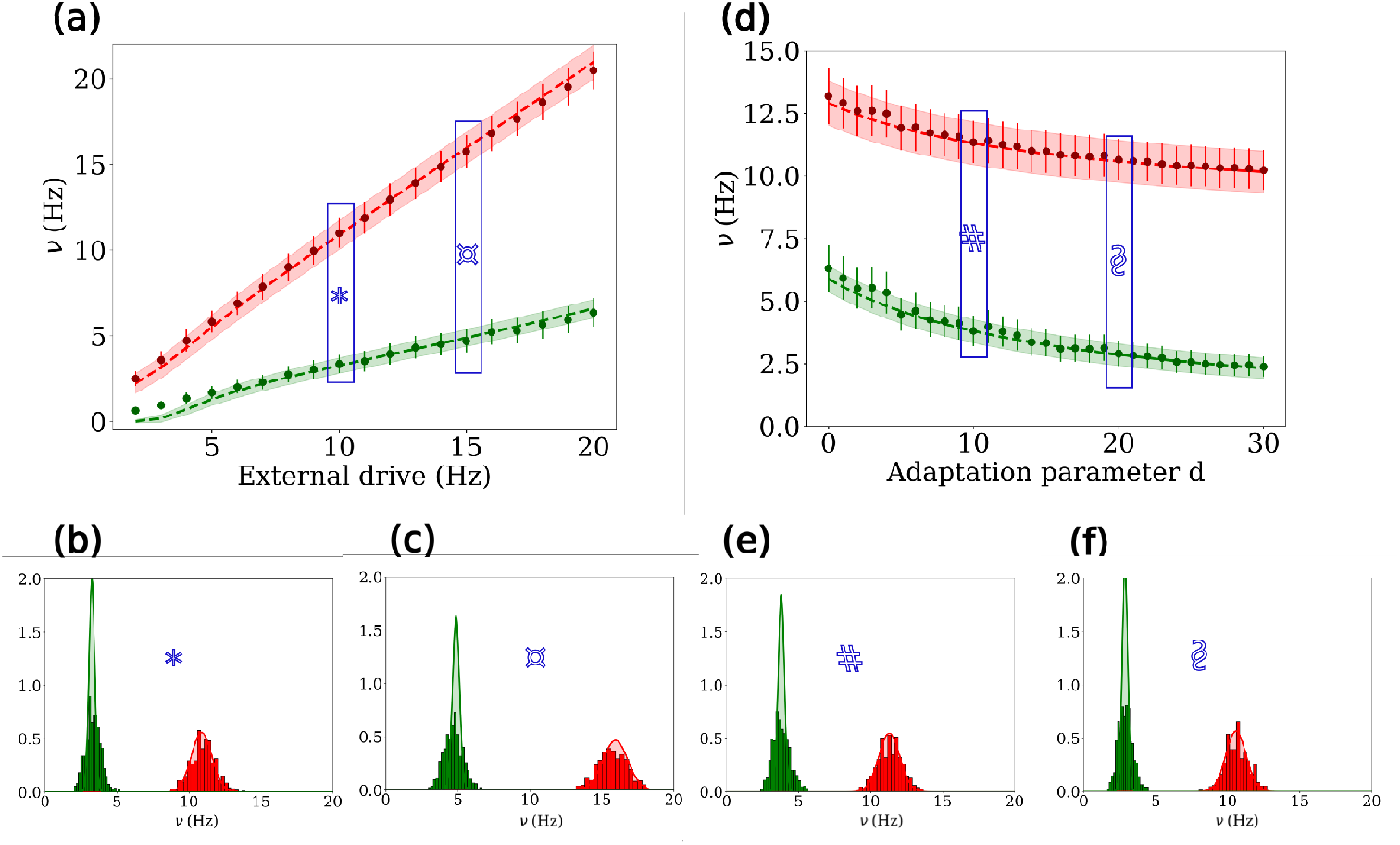
: Validation of the second order mean-field model through comparison of the mean-field prediction and the corresponding result from the spiking-network. a) Firing rate for the excitatory (green) and inhibitory (red) neurons as a function of the external drive. b-c) Firing rates distributions for two different constant inputs (indicated in (a)). d) Firing rate for the excitatory and inhibitory neurons for different values of the adaptation parameter *d*. e-f) Firing rates distribution for the two different values of *d* (indicated in (d)).

As described in the Methods section, one of the key features of the Izhikevich model is the inclusion of the slow adaptation variable. Thus, capturing the effect of the adaptation on the system is of great relevance for the validation of the mean-field. To this purpose, we show in Fig. 2.d the response of the mean-field to a constant input as a function of the adaptation parameter *d*, and in Figs. 2.e and f the firing rate distribution together with mean-field prediction for different values of *d*. As we can see, the mean-field correctly estimates the impact of the adaptation on the firing rate of the network.

### 3.2 Time-varying inputs

In the previous section, we tested our formalism for a constant external input. We now turn to study the response of the mean-field for a time-varying input. In particular, we test the mean-field response to stimuli of different amplitudes and speeds. We show in Fig. 3 the results of our simulations together with the response of the network for a Gaussian-shaped stimulus of various widths and amplitude. As we see in the figure, the mean-field model can correctly capture the response of the network to the different inputs.

**Figure 3.**
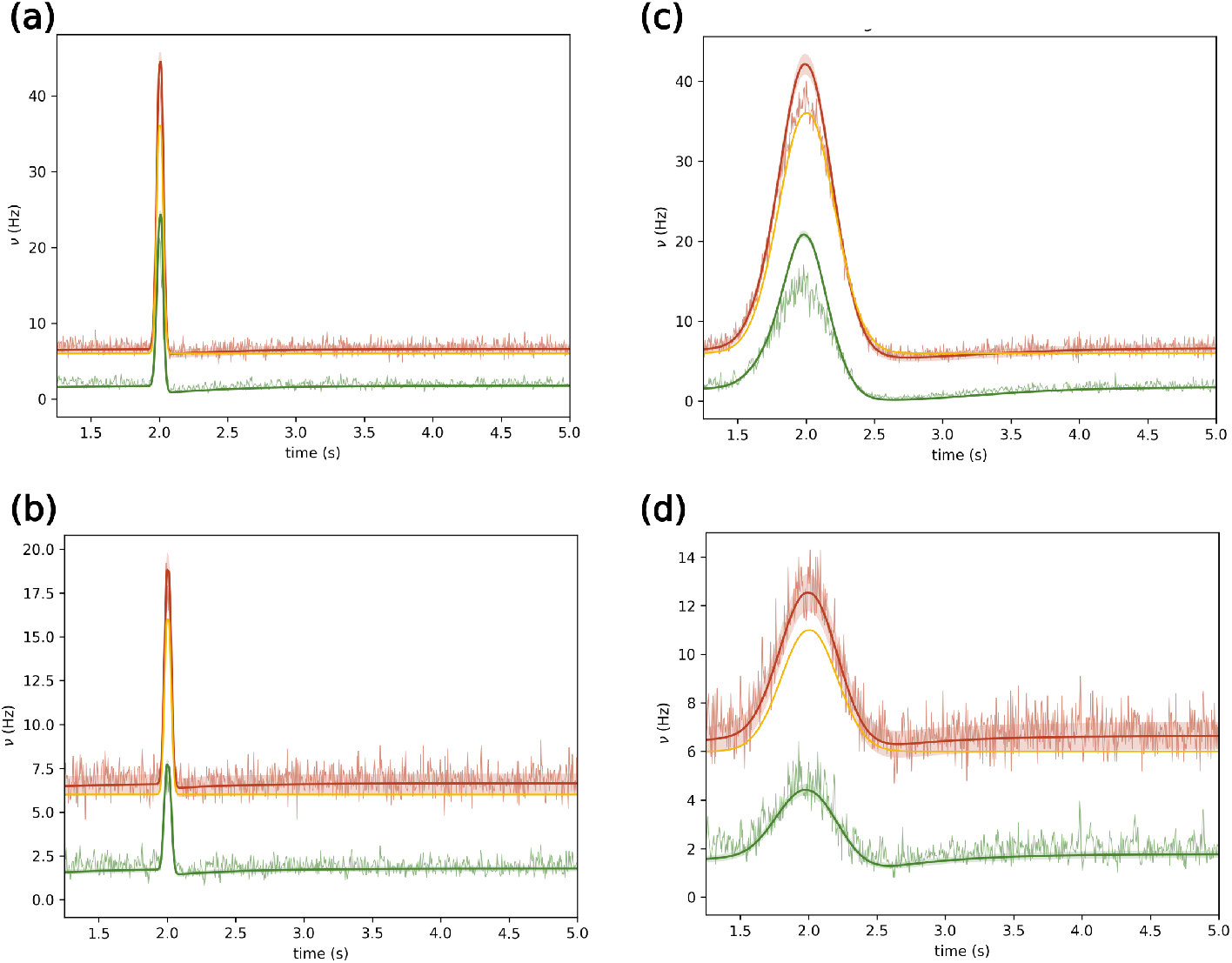
: Mean-field response to time-varying inputs. We show the response of the mean-field (darker colors) for different time-varying inputs and the corresponding network simulation (lighter colors). The applied input is shown in yellow. We see that the mean-field can correctly capture the variation in the mean-firing rates driven by fast and slow inputs of different amplitude.

### 3.3 Model robustness and parameter exploration

In principle, the estimation of the semi-analytical transfer function should be independent of the neuronal parameters, as these are explicitly taken into consideration within the formulation. This provides a large flexibility to our formalism and makes it suitable to analyze different regimes emerging from the network. To test the validity of the mean-field model under different parameters, we performed a parameter exploration and we compared the results of the network with the predictions of the mean-field. The results of this analysis are shown in Fig. 4. We show the firing rates obtained from the network and the mean-field predictions for both excitatory and inhibitory neurons, together with the adaptation variable *u*_*e*_. In particular, we explored the parameters *E*_*Iz*_ and *g*_*Iz*_ that regulate the excitability of the system. As shown in the figure, the mean-field can capture the behaviour of the network for a wide range of parameters (one order of magnitude in *g*_*Iz*_), although a discrepancy appears for low values of *E*_*I*_*z* and high *g*_*Iz*_.

**Figure 4.**
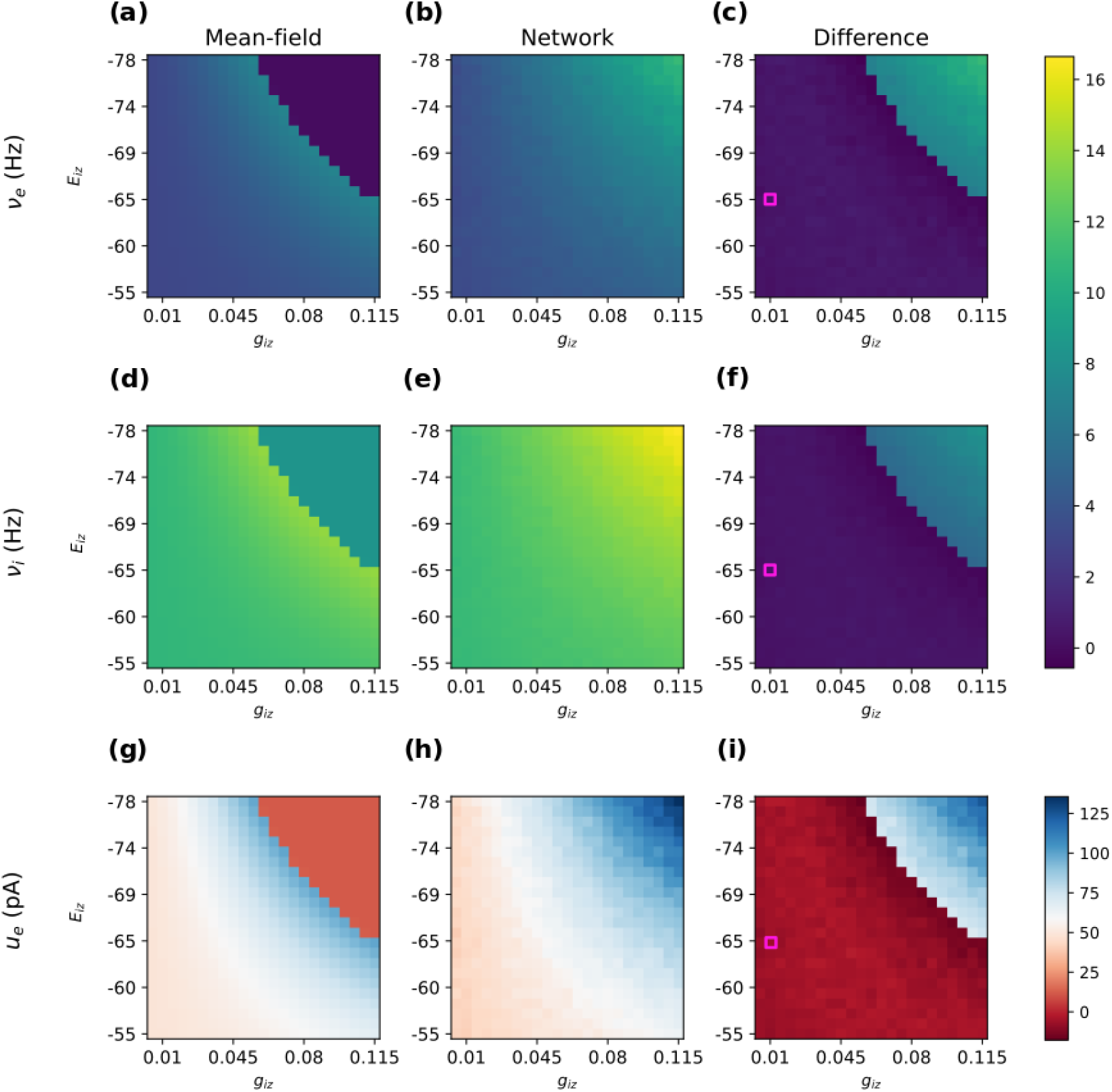
: Robustness of the mean-field model. We analyzed the validity of the mean-field for an extensive parameter exploration. The stationary values of the firing rates (a, b, c for excitatory; d, e, f for inhibitory) and the adaptation variable (g, h, i) result from the computation of the mean-field model (a, d, g) and the network model (b, e, h) as a function of the parameters *E*_*Iz*_ and *g*_*Iz*_ which regulate the excitability of the system. The original values used to build the transfer function are indicated with a pink square. We see that the mean-field remains valid for a wide range of parameters, showing the robustness of the formalism.

## 4 Discussion

In this paper, we have derived a mean-field model of populations of neurons described by the quadratic integrate-and-fire model, interacting with conductance-based synapses. We discuss this model below, its limitations and usefulness, and relate it to previous work.

A major advantage of the mean-field approximation is that it allows large populations of neurons to be studied in a computationally efficient manner, while still capturing the important features of the dynamics. By capturing the average behavior of the population, it provides a computationally efficient and mathematically tractable way to study the dynamics of large-scale neural systems. It is worth noting that our analysis is limited to simple firing patterns, where neurons fire consistently in an asynchronous-irregular pattern in response to an external stimulus, in which case the validity of our formalism is guaranteed. A more in-depth study of the capacity of this formalism to capture other dynamical behaviour of the spiking network, such as bursting, would be interesting for the future. As the model of a single neuron presents properties similar to those previously used [3, 15, 16], we can expect that dynamics such as slow-waves must exist. In this work, we focused on the basic dynamics for which this mean-field approach was designed [2] (asynchronous-irregular regimes), and we have shown that this mean-field can capture this type of dynamics for different input types. Other approaches using the same type of neuron model have been previously developed [6]. These have the advantage of not being limited in temporal resolution by the timescale of T. However, they do not account for network size effects or probability of connection (an all-to-all connection is assumed, which is far from biologically plausible).

It must be noted that a previous mean-field approach was proposed for the quadratic integrate-and-fire model [7], which was more recently extended to adaptive quadratic integrate-and-fire neurons [6]. To consider adaptation, the mean-field had to use a particular mathematical technique (called Lorentzian ansatz) to enable the inclusion of adaptation in the mean-field. In the present approach, we directly integrate adaptation in the formalism and in the transfer function [3], which is simpler. Another key difference is that these previous models work at the thermodynamic limit (when the number of neurons tends to infinity), while our approach is finite size and includes the network size in its parameters [2]. A further difference is that in the previous models [7] current-based synapses were considered, while in our formalism we adopt conductance-based synapses (also used in [6]), which are more biologically relevant. Hence, our approach can also be seen as an extension of these previous works.

Finally, it is clear that the mean-field approximation is an important tool for studying large populations of neurons. By capturing the average behavior of the population, this approach provides a computationally efficient and mathematically tractable way to study the dynamics of large-scale neural systems, up to the whole brain scale [15, 16]. Future studies could extend this approach to explore other possible dynamics, such as bursting, while also considering the limitations and advantages of other modeling approaches that use the same type of neuron model.

## 5 Conclusion

In conclusion, modelling brain activity across different scales is a complex task, and there are various methods available to achieve this. Neural mass models and mean-field approaches have emerged as powerful tools to describe the dynamics of large populations of neurons and have been successfully applied to model clinically recorded signals such as fMRI, EEG, or MEG. In this work, we have adapted a mean-field method to the adaptive quadratic integrate-and-fire neuron model, a widely used model in computational neuroscience. We have built a model of cortical columns based on a balanced spiking network and compared its dynamics to the mean-field approximation. Our results demonstrate that the mean-field approach captures the asynchronous-irregular dynamics of the spiking neural network for different input types, making it a computationally efficient and mathematically tractable way to study the behavior of large-scale neural systems.

## 6 Acknowledgment

This research resulted from a collaboration with students during the Fall School 2022 of the European Institute of Theoretical Neuroscience (www.eitn.org). This work was funded by the European Union’s Horizon 2020 Framework Program for Research and Innovation under the Specific Grant Agreement No. 945539 (Human Brain Project SGA3) and the Centre National de la Recherche Scientifique (CNRS, France).

## 7 Data Availability

The code implementation of the spiking network and mean-field simulations will be made available after publication of the work.

